# Genomic Surveillance of SARS-CoV-2 Using Long-Range PCR Primers

**DOI:** 10.1101/2023.07.10.548464

**Authors:** Sangam Kandel, Susanna L. Hartzell, Ashton K. Ingold, Grace A. Turner, Joshua L. Kennedy, David W. Ussery

## Abstract

Whole Genome Sequencing (WGS) of the SARS-CoV-2 virus is crucial in the surveillance of the COVID-19 pandemic. Several primer schemes have been developed to sequence the ∼30,000 nucleotide SARS-CoV-2 genome that use a multiplex PCR approach to amplify cDNA copies of the viral genomic RNA. Midnight primers and ARTIC V4.1 primers are the most popular primer schemes that can amplify segments of SARS-CoV-2 (400 bp and 1200 bp, respectively) tiled across the viral RNA genome. Mutations within primer binding sites and primer-primer interactions can result in amplicon dropouts and coverage bias, yielding low-quality genomes with ‘Ns’ inserted in the missing amplicon regions, causing inaccurate lineage assignments, and making it challenging to monitor lineage-specific mutations in Variants of Concern (VoCs). This study uses seven long-range PCR primers with an amplicon size of ∼4500 bp to tile across the complete SARS-CoV-2 genome. One of these regions includes the full-length S-gene by using a set of flanking primers. Using a small set of long-range primers to sequence SARS-CoV-2 genomes reduces the possibility of amplicon dropout and coverage bias.

## Introduction

Whole Genome Sequencing (WGS) is widely used for the surveillance of Severe Acute Respiratory Syndrome Coronavirus-2 (SARS-CoV-2), the causative agent of the pandemic disease COVID-19 (Wu *et al*. 2020, Zhou *et al*. 2020, Huang *et al*. 2020). At the time of writing (May 23, 2023), there are more than 15 million genomes available in the GISAID database (https://gisaid.org/) and more than 7 million genomes in GenBank (https://www.ncbi.nlm.nih.gov/sars-cov-2/). Sequencing SARS-CoV-2 genomes is crucial in tracking viral mutations that can affect viral transmission (Carabelli *et al*. 2023, Escalera *et al*. 2020, Kupferschmidt and Wadman, 2021, Brito *et al*. 2022), disease pathogenesis (Bakhshandeh *et al*. 2021), vaccine efficacy (Chatterjee *et al*. 2023, Madhi *et al*. 2021, Hoffmann et al. 2021), and virulence (Carabelli *et al*. 2023, Issa *et al*. 2020). A variety of methods, including metagenomic sequencing, hybridization capture, direct RNA sequencing, and target enrichment using multiplex PCR have been used for sequencing SARS-CoV-2 (Gerber *et al*. 2022, Liu *et al*. 2021, Rehn *et al*. 2021, Butler *et al*. 2020, Carbo *et al*. 2020, Charre *et al*. 2020, Deng *et al*. 2020, Wu *et al*. 2020, Xiao *et al*. 2020, Vacca *et al*. 2022). Most of the target enrichment methods require reverse transcription to generate a double-stranded cDNA copy of the genomic RNA (gRNA) and then utilize this cDNA as a template for DNA sequencing, using multiplex primers to cover the whole genome of SARS-CoV-2 (Grubaugh *et al*. 2019).

Target enrichment using PCR amplicons and subsequent Oxford Nanopore Sequencing is extremely popular and relatively inexpensive (∼$10 per sample), with a quick turnaround time (∼24 hours from sample to GenBank file). Target enrichment using publicly available ARTIC Network PCR primers (Tyson *et al*. 2020), Entebbe primers (1.5kb-2Kb) (Cotten *et al*. 2021), MRL primers (1.5kb-2.5kb) (Arana *et al*. 2022), and Midnight Primers (Freed *et al*. 2020) are used to sequence SARS-CoV-2 with Oxford Nanopore flow cells. Among these primer schemes, ARTIC primers and Midnight primers are the most commonly used to sequence clinical isolates of SARS-CoV-2. ARTIC primers V4 includes 98 primer pairs, each amplifying ∼400bp fragments along the viral genome, which can be sequenced on either Illumina or Oxford Nanopore platforms. The ‘Midnight primers’ have 30 primer pairs that generate amplicons with a targeted size of 1200 base pairs, taking advantage of the longer read lengths of third-generation sequencing, including Oxford Nanopore flow cells. Generation of full-length high-quality consensus sequences depends upon the quality and quantity of the viral load in clinical samples, as well as the mutations occurring within the primer binding regions of the viral genome (Kuchinski *et al*. 2021, Liu *et al*. 2021, Davis *et al*. 2021). Amplicon dropouts and coverage bias at different amplicon regions have been observed with the sequencing protocols based on ARTIC (Kuchinski *et al*. 2021, Itokawa *et al*. 2020) as well as Midnight primers (Kuchinski *et al*. 2021, Bei *et al*. 2022). Mutations within the primer binding site can prevent primer-annealing and result in ‘dropout’ or loss of that amplicon, leading to incomplete genome sequences (Bei *et al*. 2022, Sanderson and Barret 2021). Furthermore, primer-primer interactions could result in amplification bias of interacting amplicons (Itokawa *et al*. 2020), resulting in coverage bias and affecting the identification of mutations in the viral genome that are key in the nomenclature of emerging variants.

The variants of SARS-CoV-2 are determined by a combination of several mutations that occur mainly within the Spike gene. For example, in the Alpha variant (B.1.1.7), there are 14 critical lineage-defining mutations within the S gene (Galloway *et al*. 2021). Similarly, Omicron subvariant B.1.1.529 has 60 mutations within the viral genome, including 15 key mutations within the receptor binding domain (He *et al*. 2021). The characteristic mutation within the S gene for the Alpha variant B.1.1.7 (Clark *et al*. 2021, Meng et al. 2021) and the Omicron variants B.1.1.529, BA.1, BA.1.1 (Clark *et al*. 2021) is the deletion of two amino acids at positions 69 and 70 (del H69/V70) (https://covariants.org). This deletion inhibits the PCR amplification of the S-gene (S-Gene Target Failure, or SGTF) in diagnostic PCR assays such as the ThermoFisher TaqPath™ COVID-19 Combo Kit RT-PCR (Clark *et al*. 2021, Davies *et al*. 2021) that targets the N, ORF1ab, and S gene regions. This deletion (del H69/V70) results in a false-negative result for the S-gene targeted diagnostic test. SGTF became a proxy for early detection of Alpha and Omicron B.1.1.529 variants (Galloway *et al*. 2021). In addition, a mutation at position 27,807 (Cytosine substituted to Thymine) within amplicon 28, also a primer annealing site (Primer 28_LEFT, pool B of Midnight primer) (Supplementary figure 1, IGV plot), caused a common dropout in the Delta variant genome when using Midnight Primers (Kuchinski *et al*. 2021). Spiking Primer pool B with a custom primer designed by substituting Cytosine with Thymine base not only corrected the dropout but also increased the coverage at this region (Constantinides *et al*. 2022). Furthermore, the genome sequences of two BA.2 Omicron variants from Arkansas (GenBank Accession: OM863926, ON831693) sequenced using Midnight Primers in Oxford Nanopore GridION have a complete dropout at amplicon region 21 (20,677-21,562). The Omicron and the Alpha variant waves taught us that tests and primers designed towards regions within the S gene could result in false-negative tests because this gene encodes a surface protein, subjecting it to varying selectional pressures (Julenius and Pedersen 2006). Variations can lead to problems that are troublesome in deciding the public health interventions needed to control the transmission and spread of COVID-19 disease.

Multiplex primers used to sequence SARS-CoV-2 viral isolates must be targeted to bind regions that are conserved with little variance to avoid dropout failures secondary to the primers not binding. Long-range PCR primers targeting the amplification of 4500bp can prevent the ‘S-gene dropouts’, as the primer binding sites flanking the S-gene region are located within highly conserved regions on either side of the S gene. The S gene is approximately 3,822 base pairs long and stretches between the nucleotide position 21,563 to 25,384 along the viral genome. Therefore, these long-range PCR primers can generate amplicons around 4500bp that will cover the entire S gene, making the chances of amplicon dropout within the S-gene minimal. We have previously demonstrated whole-genome cDNA sequences from Mumps genomes using long-range PCR yielding fragments of ∼ 5000 bp in length from buccal samples (Alkam *et al*. 2019). Through our work in SARS-CoV-2, we have identified conserved regions that flank the S gene (Wassenaar *et al*. 2022). In this study, we designed long-range PCR primers to target these conserved S gene areas and sequence SARS-CoV-2 isolates. Our objective was to improve the quality of the sequences generated and minimize the amplicon dropouts, as the designed primers are outside the highly variable regions.

## Results

Long-range primers were used to sequence four samples identified as: V05476 _11.6, V05450 _15.1, V06110 _14.3, V06106 _18.3 with cycle threshold (CT) values of 11.6, 15.1, 14.3, and 18. 3 respectively on an Oxford Nanopore GridION machine. A total of 4.8 million reads were generated from 4 samples with N50 of 2,640 bases after 28 hours of sequencing. The mean read coverage was approximately the same (7529, 7646, 7673, and 7725, respectively) for the four samples (Table 2). All the samples had high genome coverage (>98%; see Figure 2), and each was assigned the BA.5 variant of Omicron. The number of reads mapped to each amplicon position is summarized in Figure 3 and Table 3. Out of seven amplicons, amplicon 4 had the highest number of reads mapped to the reference.

**Figure 1:**
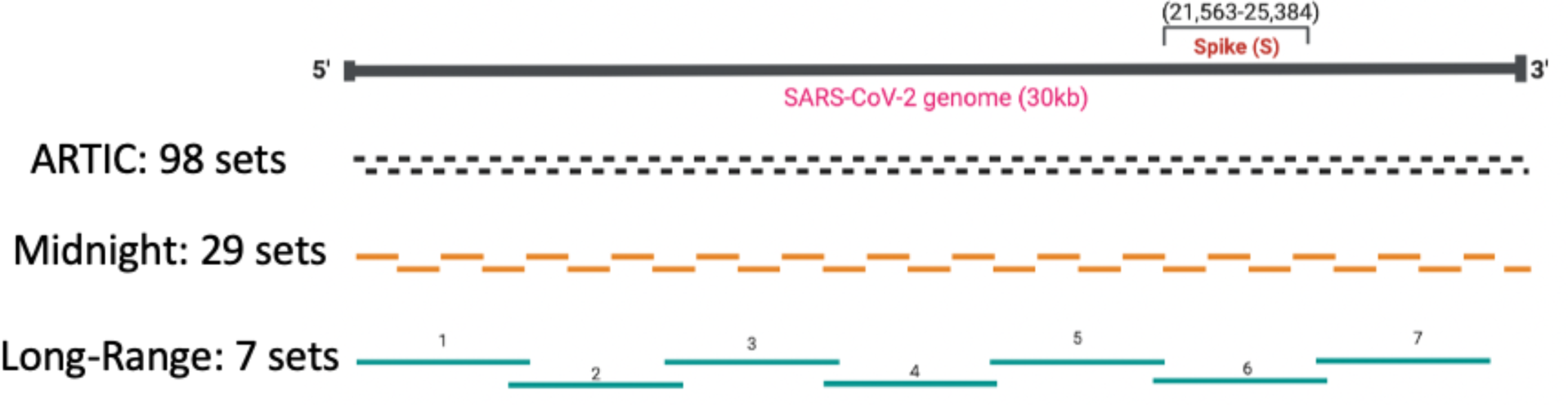
Comparison of ARTIC, Midnight, and Long-Range PCR primers. Long-range PCR primers used in this study include seven primer pairs to sequence the whole genome of SARS-CoV-2. The entire S-gene is sequenced by just one long-range primer. The horizontal dotted lines represent the viral genome segments amplified by each primer set.

**Figure 2:**
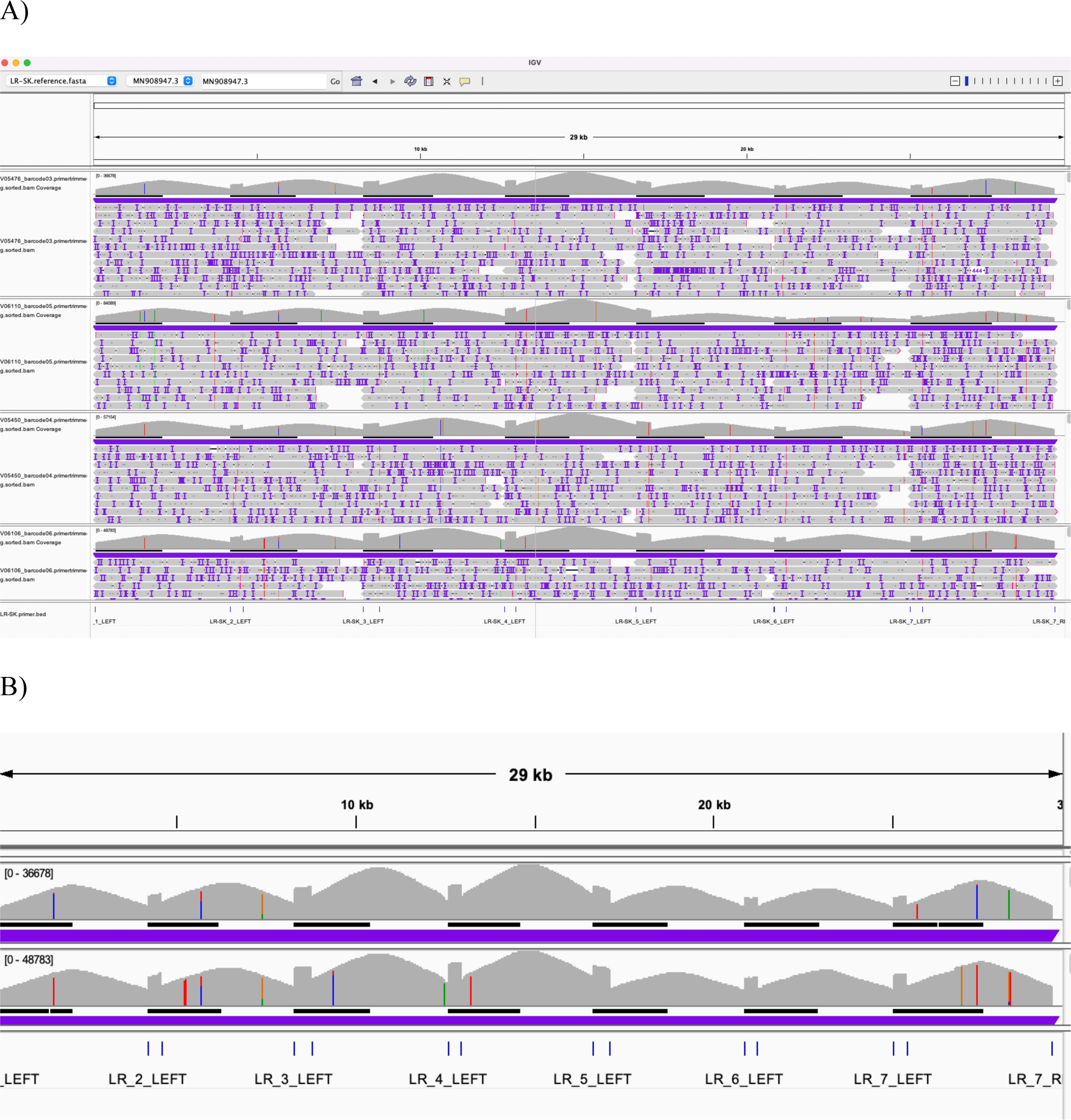
IGV plot showing seven different amplicons mapped to the SARS-CoV-2 reference genome for four samples with low CT values. A) Samples with CT values 11.6, 15.1,14.3, and 18.3 from top to bottom, respectively. B) IGV plot for two samples (zoomed for the sample with CT values of 11.6 and 18.3 from top to bottom, respectively. The scale [0-36678] for the top and [0-48783], respectively, represents the range of the total number of the quality filtered reads that mapped to each amplicon region. The details of the reads mapped to different amplicon regions for four samples sequenced are summarized in Table 3 and Figure 3.

**Figure 3:**
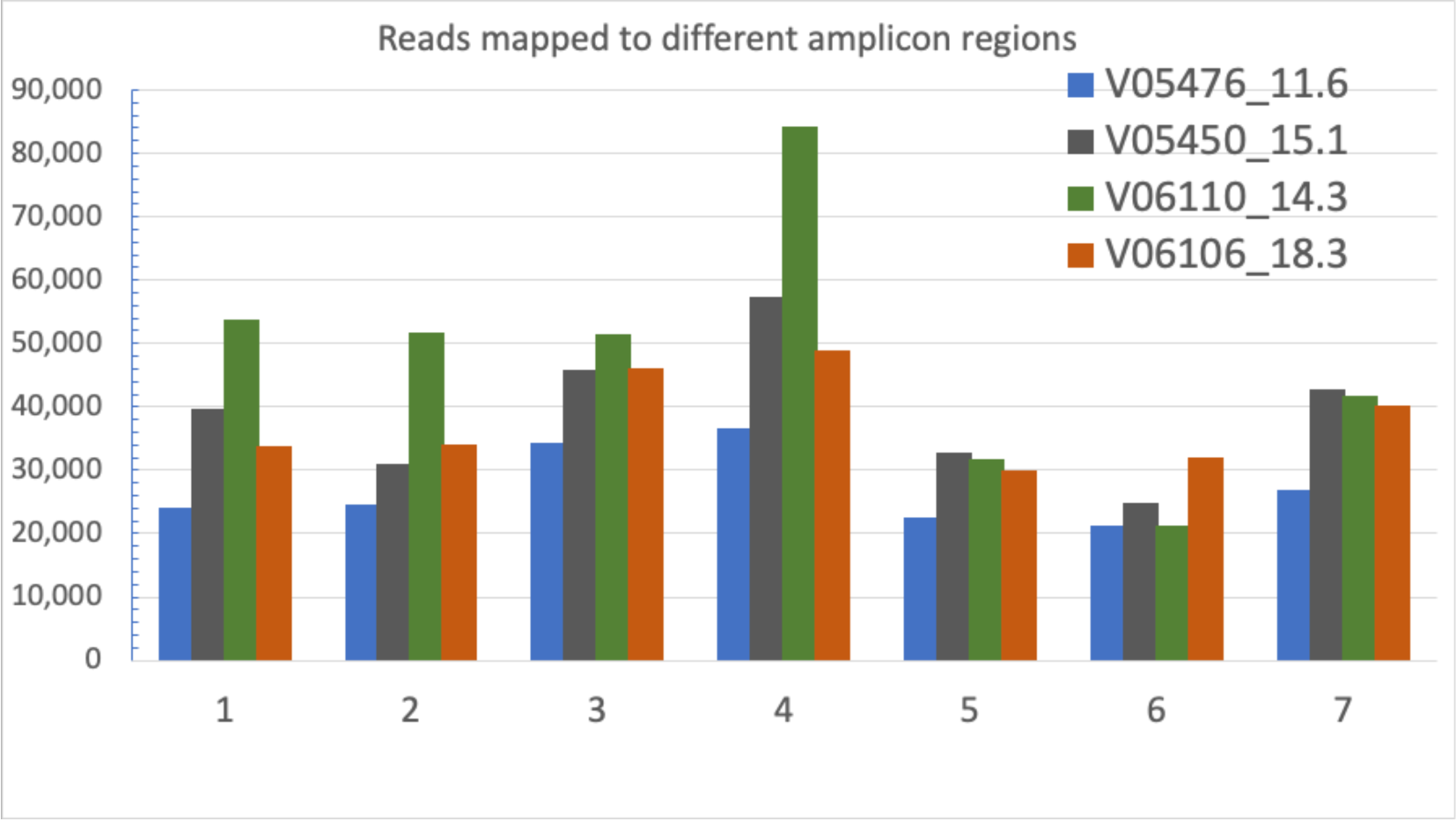
Histogram showing the number of reads mapped to each amplicon in the primer scheme. The ‘align_trim’ report file from the ARTIC pipeline was used in the ONTdeCIPHER tool. If the alignment length between the read and reference is <0.95% of the amplicon length, the read is discarded from the coverage plot. Amplicons are marked as dropped out if the total number of reads assigned to an amplicon is below 50. (X-axis: Seven amplicon regions tiled across the whole genome of SARS-CoV-2, amplifying seven regions along the viral genome. Y-axis: Total number of sequencing reads mapped to each amplicon region).

**Table 1:** Comparison of ARTIC, Midnight, and Long-range primers used to sequence SARS-CoV-2 clinical isolates

|  | ARTIC | Midnight | Lon-range |
| --- | --- | --- | --- |
| Number of primers sets | 98 | 30 | 7 |
| Amplicon size (base pairs) | 400 | 1,200 | 4,500 |

**Table 2:** Sequencing summary of four samples showing different quality metrics.

| Sample | Raw reads | Filtered reads | Mapped to reference | Mean read coverage | Variant |
| --- | --- | --- | --- | --- | --- |
| V05476_11.6 | 646,100 | 263,335 | 98.4 % | 7,529 X | BA.5.1 |
| V05450_15.1 | 817,300 | 373,648 | 98.4 % | 7,646 X | BA.5.2.1 |
| V06110_14.3 | 974,000 | 452,910 | 98.4 % | 7,672 X | BA.5.3.1 |
| V06106_18.3 | 759,000 | 355,864 | 98.4 % | 7,725 X | BA.5.2.1 |

**Table 3:** Total number of raw reads, filtered reads, and reads that mapped to the reference genome at seven amplicon regions.

| Sample | V05476_11.6 | V05450_15.1 | V06110_14.3 | V06106_18.3 |
| --- | --- | --- | --- | --- |
| Amplicon 1 | 24006 | 39741 | 53725 | 33702 |
| Amplicon 2 | 24656 | 30882 | 51645 | 34150 |
| Amplicon 3 | 34223 | 45931 | 51497 | 46092 |
| Amplicon 4 | 36750 | 57256 | 84273 | 48869 |
| Amplicon 5 | 22471 | 32822 | 31826 | 29876 |
| Amplicon 6 | 21170 | 24926 | 21167 | 31987 |
| Amplicon 7 | 26924 | 42728 | 41629 | 40237 |
| Total mapped | 190,200 | 274,286 | 335,762 | 264,913 |
| Total raw reads | 646.1 K | 817.3 K | 974 K | 759.7 K |
| Filtered reads | 263,335 | 373648 | 452910 | 355864 |

A 96-well plate containing samples with different CT values spanning from 11 to 16 (n=19), 17 to 20 (n=14), 21 to 25 (n=15), 26 to 30 (n=15), 31 to 35 (n=15), and 36 to 42 (n=16) were sequenced using long-range and Midnight primers for comparison. With long-range primers, 100% of the samples with CT values 11 to 16 passed quality, whereas 95 % of samples within the range of this CT value passed quality when sequenced with midnight primers. Long-range primers were as good as midnight primers for sequencing samples with CT values between 17-20 (Long-range: 73% and Midnight: 88% passing quality). For samples with CT values of 21-25, 47% passed quality with Midnight primers, whereas 33% passed quality with long-range primers. With midnight primers, only two samples passed quality with CT values greater than 26. The long-range and the midnight primers generated no quality sequences in those samples with CT values greater than 26 (Figures 4, 5, 6, and Table 4).

**Figure 4:**
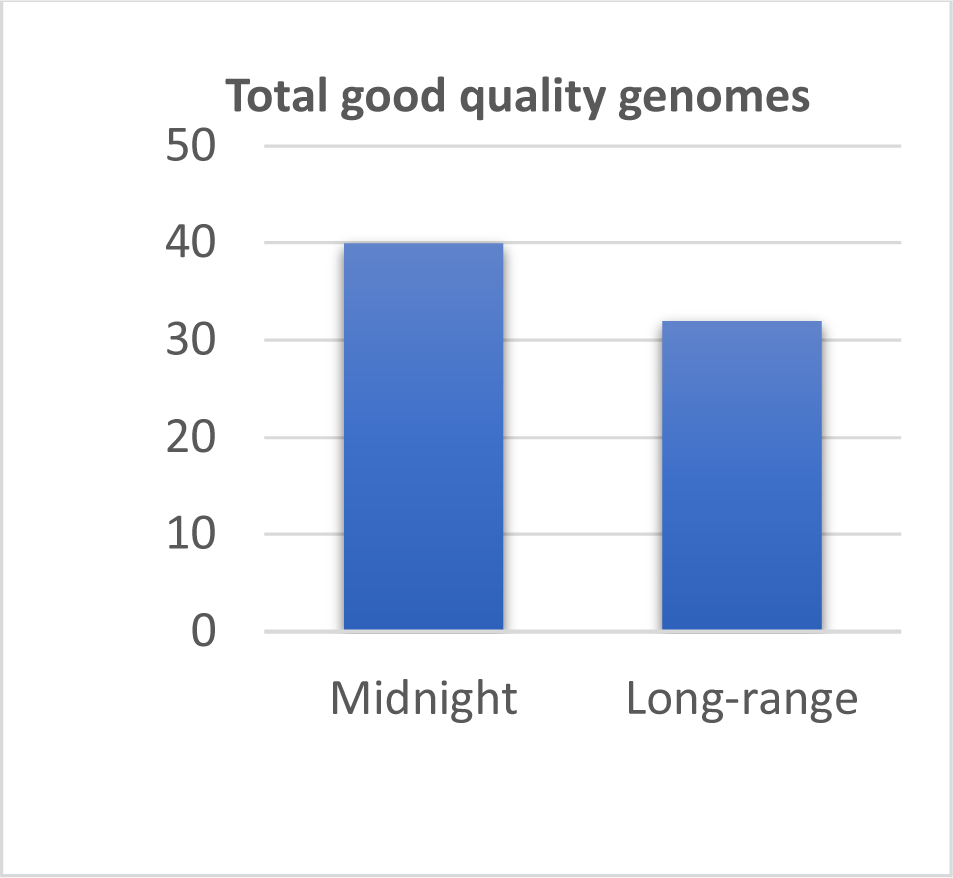
Bar chart showing the total number of samples passing quality. A 96-well plate with samples of different CT values was sequenced using Long-range and Midnight primers for comparison. Long-range primers and midnight primers work to accurately assign lineages and generate good-quality genomes for GenBank.

**Figure 5:**
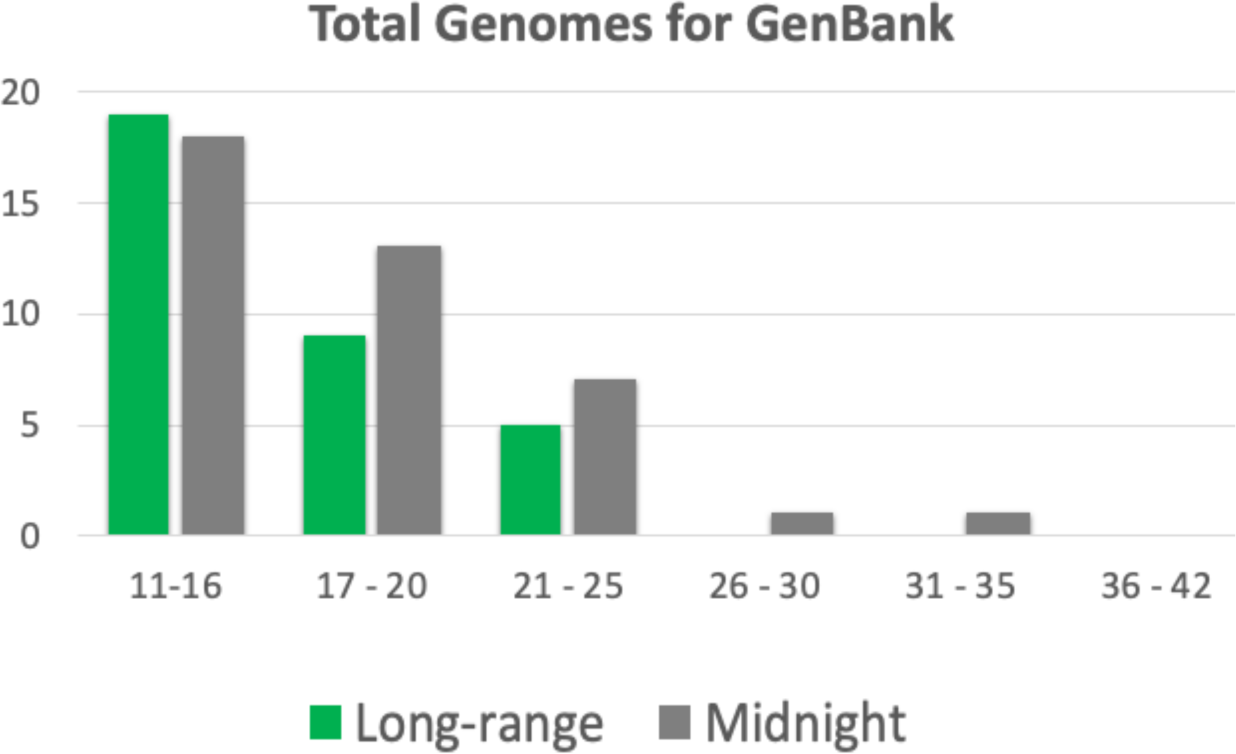
Bar chart showing samples sequenced with Midnight and Long-range primers with different CT values that passed quality. X-axis: CT value range, Y-axis Number of genomes passing quality.

**Figure 6:**
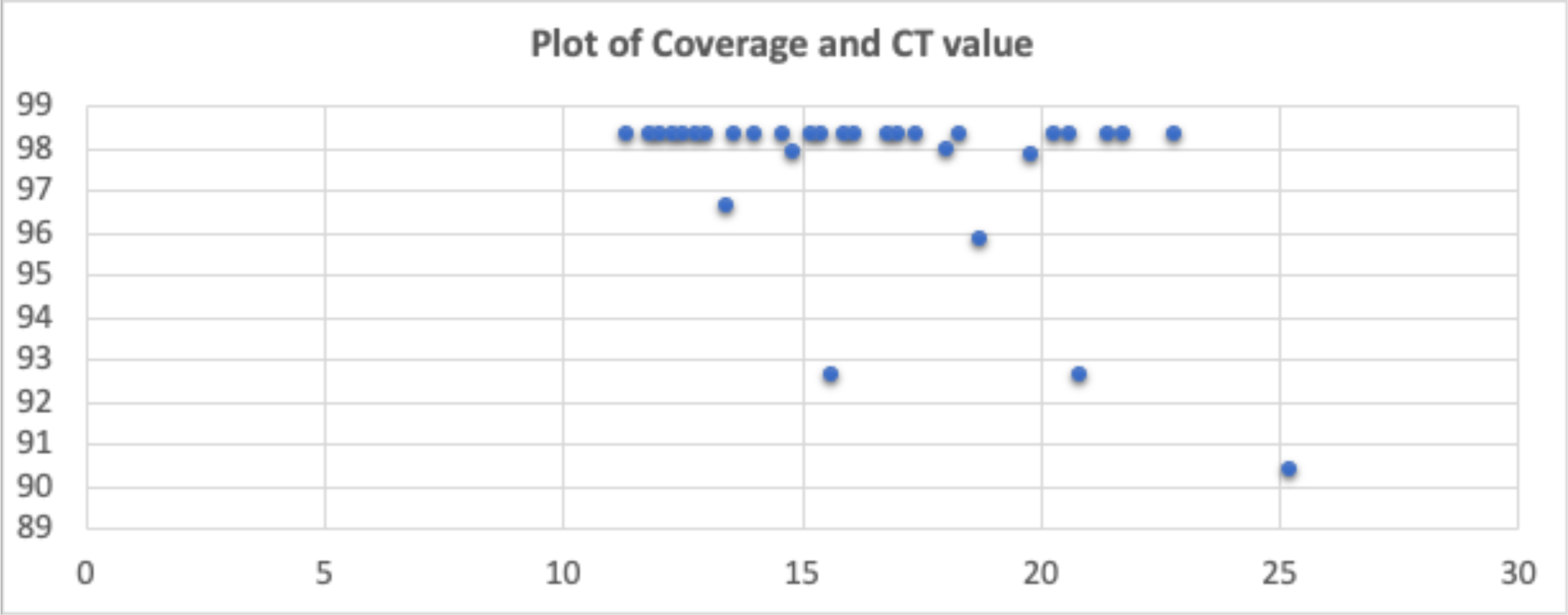
Plot of genome coverage and CT values for the genomes that passed quality when sequenced using long-range primers. Long-range primers are effective in sequencing samples with CT values less than 20 to get at least 99 % genome coverage.

**Figure 7:**
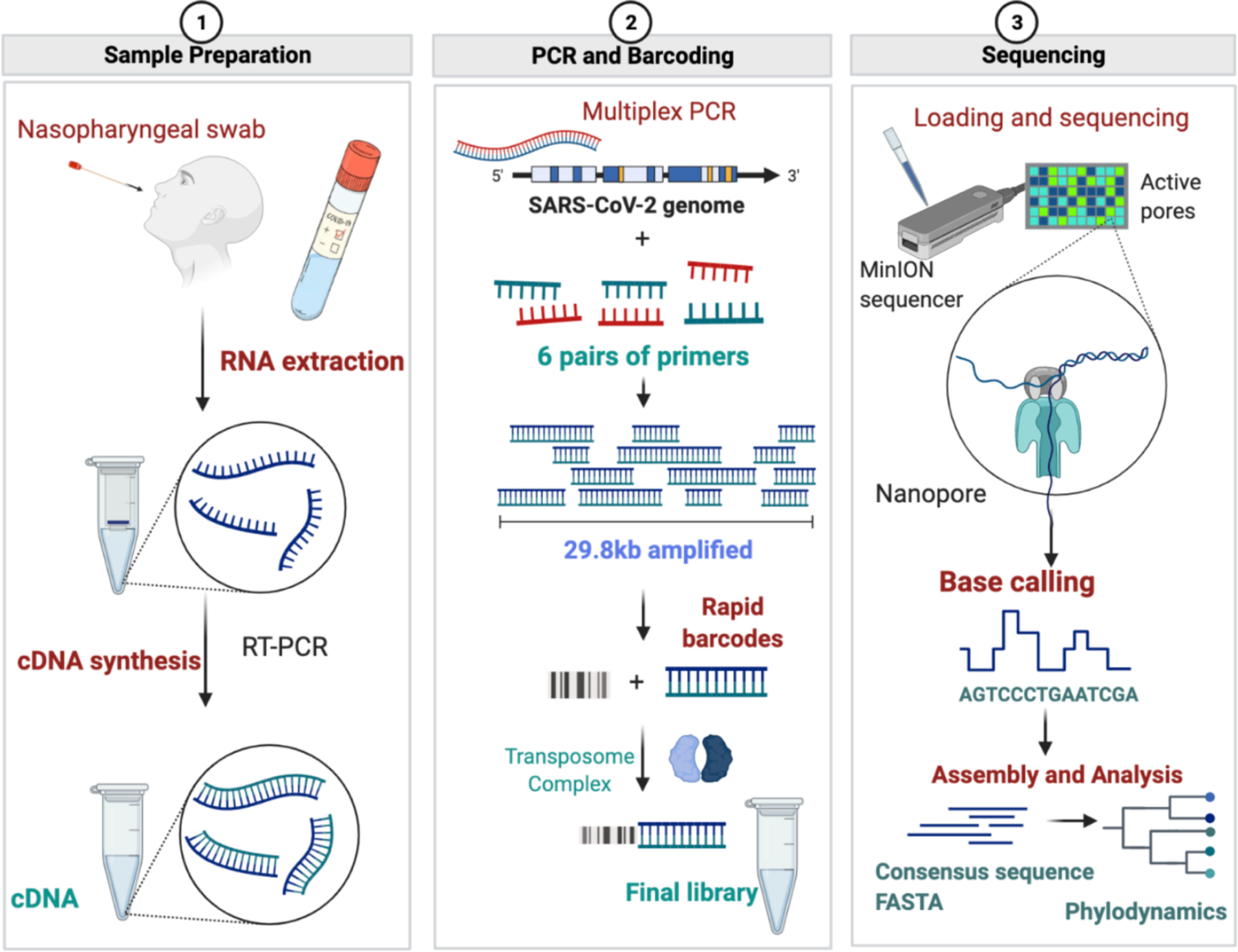
Diagrammatic representation of Oxford Nanopore Sequencing of SARS-CoV-2 using long-range PCR primers. (Figures made using BioRender.com)

**Table 4:** Comparison of total samples passing quality standards by CT values.

| CT values | Samples | GenBank (Long-range) | GenBank (Midnight) |
| --- | --- | --- | --- |
| 11 - 16 | 19 | 19 (100%) | 18 (95%) |
| 17 - 20 | 14 | 9 (73%) | 13 (88%) |
| 21 - 25 | 15 | 5 (33%) | 7 (47%) |
| 26 - 30 | 15 | 0 | 1 (0.07 %) |
| 31 - 35 | 15 | 0 | 1 (0.07%) |
| 36 - 42 | 16 | 0 | 0 |
| Total | 94 | 32 | 40 |

Although the samples from a 96-plex sequencing run that passed quality were accurately assigned to a lineage, we have found, in some cases, there was low coverage of some regions. For this reason, we developed alternative primers to address the low coverage of these amplicons (Supplementary Figure 3) and to target the recent Omicron variant XBB. With optimized PCR conditions, this alternative primer generated high-quality genomes with a lineage assigned to the consensus sequence of the genome (Supplementary Table 1, Supplementary Figure 4). As the virus continues to mutate, it will likely be necessary to adjust the primers to maintain optimal coverage for all regions.

## Discussion

We have developed and evaluated novel long-range primers to sequence SARS-CoV-2 clinical isolates using Oxford Nanopore sequencing. These novel primers can amplify regions of ∼4,500 base pairs. Using our primer set, the entire S-gene was sequenced using a single primer set. We compared the performance of long-range primers with midnight primers and found that long-range primers work as good as the midnight primers regarding the quality of genome sequences and coverage. This finding depends upon the amount of viral RNA in the sample.

We used 7,000 reference genomes from GISAID to generate a consensus sequence to design these long-range primers. Genome coverage is improved when primer schemes are created using multiple reference genome sequences compared to those designed using a single reference genome (Bei *et al*. 2022). ARTIC v3 and Midnight-1200 primers were designed using just one reference genome of SARS-CoV-2. In contrast, other primer schemes, such as the updated ARTIC (ARTIC v4.1), VarSkip Short v2, and VarSkip Long primers, were designed using multiple reference genomes. Long-range PCR primers can minimize the amplicon dropout due to mutations within the primer binding site (Bei *et al*. 2022).

After the ARTIC protocol was made public on January 22, 2020, these primers were adopted globally to sequence millions of SARS-CoV-2 genomes. After the introduction, there have been several improvements and updates to these primers to resolve dropouts and improve sequencing coverage (Grubaugh *et al*. 2019, Tyson *et al*. 2020, Davis et al. 2020). In addition to ARTIC primers, midnight primers that are extremely popular for sequencing SARS-CoV-2 clinical isolates using Nanopore sequencing were also updated to resolve amplicon dropouts and coverage bias along different regions of the viral genome (Constantinides *et al*. 2021). Several studies have been conducted to compare different sequencing protocols, using multiplex PCR primers to increase the genome coverage, improve the sequencing reading quality, eliminate amplicon dropouts, and improve coverage bias at different amplicon regions (Lambisia et al. 2022, Constantinides *et al*. 2022, Bei *et al*. 2022). As the virus mutates and spreads throughout communities, the primers and protocols need to be updated to avoid amplicon dropouts and avoid coverage bias.

Long-range primers to sequence SARS-CoV-2 have not been developed apart from a few primer schemes amplifying regions up to 2,500 base pairs (Arana *et al*. 2022). Because the S-gene is approximately 3,821 base pairs, amplifying the entire S-gene requires more than one primer. Therefore, mutations within S-gene could result in dropout within S-gene. As an alternative to this problem, leveraging the long-read sequencing available with Oxford Nanopore flow cells, we have developed long-range primers, which sequence the entire S-gene using just one primer pair, thereby eliminating the possibility of amplicon dropout due to mutations within S-gene.

A limitation of this approach is that a mutation within the primer binding sites can result in a drop out of that entire region, leading to a more significant gap in the consensus sequence that significantly affects the quality of the genome sequence. However, since the primer sites were designed using conserved regions, we anticipate that this will continue to work, although, as necessary, it is easy to update the primers for novel strains. Another limitation is associated with viral load in the sample. We have found that although these long-range primers can amplify larger segments of the viral genome, these primers are not well suited to sequence samples with higher CT values (greater than 25).

Although WHO lifted the global health emergency due to a significant reduction in positive cases, we are entering into a new phase of COVID-19 as 1 out of 10 people have long-haul COVID (Thaweethai *et al*. 2023). Looking back to historical epidemics due to coronavirus and the evolutionary relatedness of the SARS-CoV-2 with previous outbreaks of SARS and MERS, future pandemics are inevitable. COVID-19 is still circulating as local outbreaks continue. The long-range PCR method outlined here can help with surveillance of community infections through wastewater monitoring. With single reads over the entire S-gene region, it is possible to quantitate variant diversity within a sample. This will allow monitoring of emerging variants as well as keeping track of known variants of concern.

## Methods

### Primer design

A total of 7,046 Omicron sub-variants (BA.2, BA.3, BA.4, BF.5, BA.5.1, BA.5.2.1, BA.5.2) genomes were downloaded from GISAID on August 12, 2022. Pangolin v4.0.6 (O’Toole *et al*. 2021) was used to assign lineages to the genomes, and any ‘unclassified’ genomes were removed. Genome sequences that were 100 % identical were then filtered out to avoid redundancy, and genome sequences having gaps of 5Ns or more in their sequences were removed that resulted in 1,205 high-quality genomes that were used for multiple sequence alignment using MAFT (Katoh *et al*. 2019). MSA Viewer (https://www.ncbi.nlm.nih.gov/projects/msaviewer/) was used to visualize the alignment, and consensus sequences were downloaded from MSA Viewer. PrimalScheme (Quick *et al*. 2017) was used to generate primer schemes using the consensus genome generated from the alignment of 1205 high-quality genomes, including different sub-variants of Omicron. Primers were designed using the PrimalScheme tool using the command line: primalscheme multiplex <fasta-file> –a 4500 –o <path-to-output> –n <primers_name= –t 30 –p –g Primers were ordered from Integrated DNA Technology (IDT) (Coralville, IA) in lab-ready form. Individual primers in each pool were mixed and resuspended to a final concentration of 100 µM. Each primer was normalized to 3 nmol during synthesis. Primers were diluted in Nuclease-free water (Sigma) to use in a final concentration of 10 µM.

High-quality genomes were downloaded from GenBank, and a consensus sequence was generated using the most recent dominant variants of SARS-CoV-2 from GenBank collected between December 2022 and March 2023. Quality filtering was done to include only those genomes that did not contain any non-ATCGN bases and those that did not have any ‘N’s in the genome sequence. The consensus sequence from this set of genomes was used to manually design the alternative primers, including amplicons 2, 3, 5, 6, and 7.

### In-vitro validation of primers

MFEprimer tool was used to predict the various quality metrics of the primer scheme designed using PrimalScheme. Primers 5_LEFT and 7_LEFT were predicted to form bases complementarity at the 3’ ends at five bases (Supplementary figure 2). Since these primers do not interact with each other, this did not affect the coverage (Figure 2: IGV plot of 4 samples).

### Detection and quantification of SARS-CoV-2 viral mRNA

All the samples used in this study were collected at Arkansas Children’s Hospital and the University of Arkansas for Medical Sciences as routine surveillance between (November 2022 – Jan 2023). Nasal swab samples were collected in a 3 mL M4RT transport media (Remel, San Diego, CA). Samples were tested for the SARS-CoV-2 using the Aptima® SARS-CoV-2 (Panther® System, Hologic, San Diego, CA) nucleic acid amplification assay. Positive samples were stored frozen at –80°C until they could be further processed.

### RNA extraction, Library Preparation, and Whole genome sequencing

Two hundred fifty microliters of viral transport media from clinical nasal swabs were used for viral RNA extraction using the MagMax Viral/Pathogen Nucleic Isolation Kit (Applied Biosystems) on the Kingfisher Flex automated instrument (Thermofisher). Viral RNA was reverse transcribed using LunaScript RT SuperMix (NEB #E3010) to generate cDNA as described (Freed *et al*. 2020). Each reverse transcription reaction contained 8 μL template RNA and 2 μL LunaScript RT SuperMix (NEB #E3010). The reaction condition for reverse transcription was: 25 °C for 10 min, followed by 50 °C for 10 min and 85 °C for 5 min. Subsequent cDNA amplification and sequencing were done using a modified Midnight protocol. In brief, viral cDNA was used in the tiling PCR method to amplify the SARS-CoV-2 viral genome using long-range PCR primers in 2 reaction pools. These primers generate PCR amplicons of around 4,500 bp size. Pool A consisted of the primers specific to amplicon regions 1, 3, 5, and 7, whereas Pool B consisted of the primers specific to amplicon regions 2, 4, and 6. A 25 µL PCR reaction mixture contained 2.5 µL template cDNA, 8.9 µL RNase-free water, 1.1 µL Primer pool A or Primer pool B (10 µM), 12.5 µl Q5 Hot Start HF 2x Master Mix (NEB # M0494X). The PCR conditions used were: 98 °C for 30 seconds (Initial denaturation), 40 cycles of: 98 °C for 10 seconds (Denaturation), 65 °C for 30 seconds followed by 72 °C for 5 minutes (Annealing and extension), and a final extension of 72°C for 5 minutes. Pool 1 and Pool 2 amplicons were pooled together, and 7.5 µL of each sample were barcoded using 2.5 µL of rapid barcodes available with the kit SQK-RBK004 (ONT). Barcoded samples were pooled together and cleaned using 0.8 X AMPure beads (Beckman Coulter, USA) to retain larger DNA fragments. The sequencing library was prepared using sequencing kit SQK-RBK004 (ONT), loaded onto a MinION flow cell (ONT), and sequenced for 28 hours using a Minion R9.4.1 flow cell on GridION with the MinKNOW application.

### Bioinformatics analysis

Basecalling and demultiplexing the sequencing reads in FAST5 format was done in real-time using Guppy v5.0.7 (Wick *et al*. 2019) with a high-accuracy model. A minimum quality score of 9 was used to remove low-quality bases. Demultiplexed FASTQ files were processed using the ARTIC Network Bioinformatics pipeline (https://artic.network/ncov-2019/ncov2019-bioinformatics-sop.html). Sequencing reads were quality filtered using artic gupplyplex method, and reference-based genome assembly was done using medaka from the artic minion method of the ARTIC bioinformatics pipeline. ONTdeCIPHER (Cherif *et al*. 2022) was used for generating visualization plots for genome coverage at different amplicon regions. The consensus sequence was generated by mapping to NC_045512.2 as a reference. Read depth was calculated using samtools depth (Li *et al*. 2009). Pangolin v4.0.6 was used to assign lineages to the genomes sequenced (O’Toole *et al*. 2021). Nextclade (Aksamentov *et al*. 2020) was used for assigning lineage as well as visualization and comparison of mutations within the viral genome.

## Data availability

The samples used in this study were sequenced for SARS-CoV-2 variant surveillance at Arkansas Children’s Hospital and Arkansas Children’s Research Institute. They were sequenced on either Nanopore GridION machine with the Midnight primers or on the Illumina NextSeq using ARTIC v.4 primers. The samples and their GenBank accession numbers are summarized in Table 5.

**Table 5:**
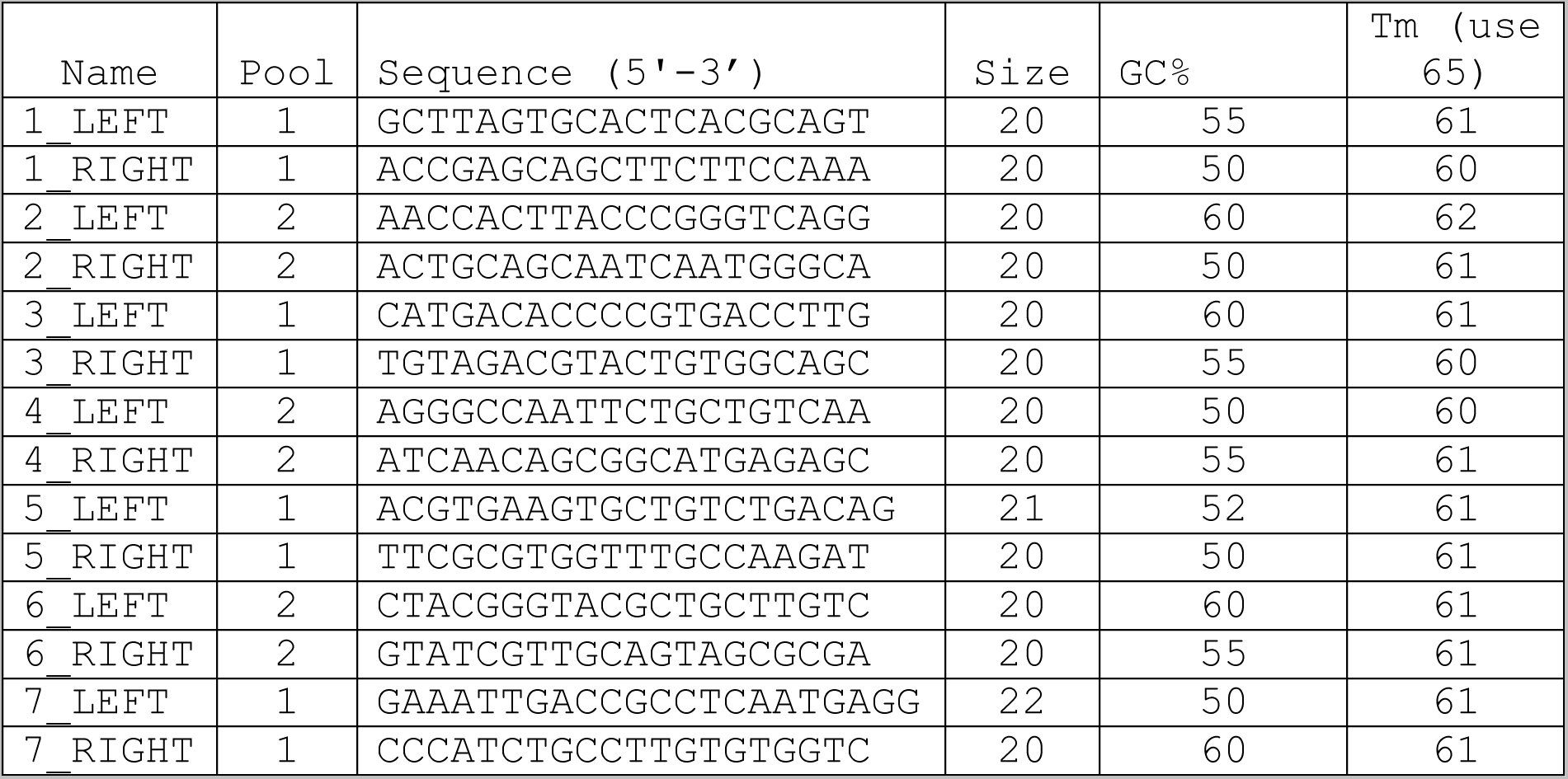
List of 7 primer pairs designed using PrimalScheme.

**Table 6:** Optimized PCR conditions for cDNA amplification to sequence SARS-CoV-2 clinical isolates.

| Steps | Temperature | Time | Cycles |
| --- | --- | --- | --- |
| Initial denaturation | 98°C | 30 seconds | 1 |
| Denaturation | 98°C | 10 seconds | 40 |
| Annealing and extension | 65°C | 30 seconds |  |
|  | 72 °C | 5 minutes |  |
| Final Extension | 72°C | 5 minutes | 1 |
| Hold | 4°C | ∞ |  |

**Table 7:**
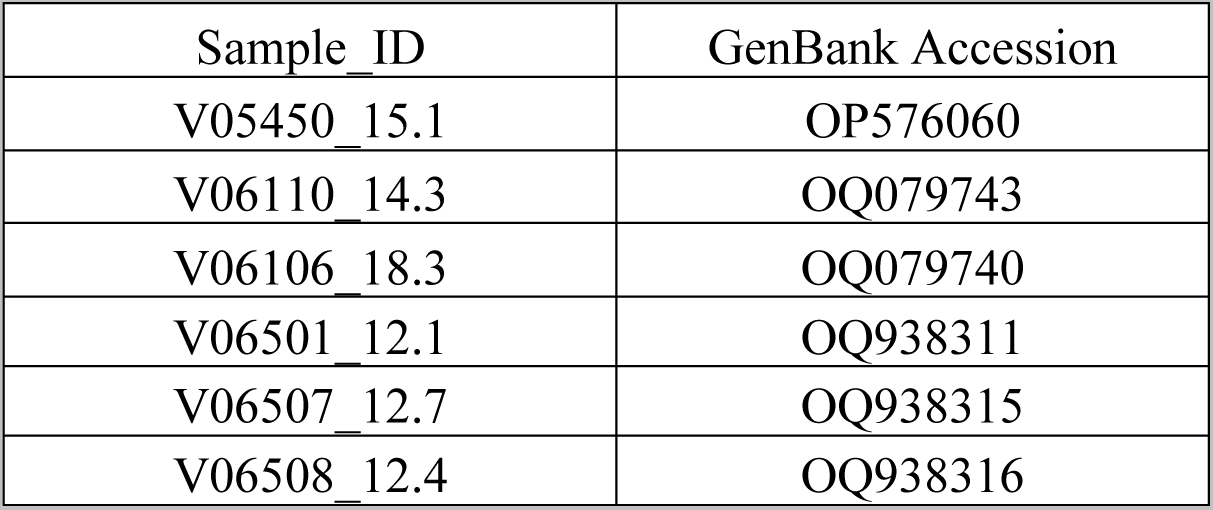
GenBank accession number of the samples used to validate this study’s long-range primers.

| Sample_ID | GenBank Accession |
| --- | --- |
| V05450_15.1 | OP576060 |
| V06110_14.3 | OQ079743 |
| V06106_18.3 | OQ079740 |
| V06501_12.1 | OQ938311 |
| V06507_12.7 | OQ938315 |
| V06508_12.4 | OQ938316 |

## Author Contributions

SK, DWU, and JLK designed the project. AKI, SLH, GAT, JLK processed the sample and did RNA extraction. SK did the sequencing, analyzed data, and wrote the first draft of the manuscript. DU and JLK supervised the project and participated in data analysis and manuscript preparation. All authors approved the submitted version.

---

**Supplementary Figure 1:**
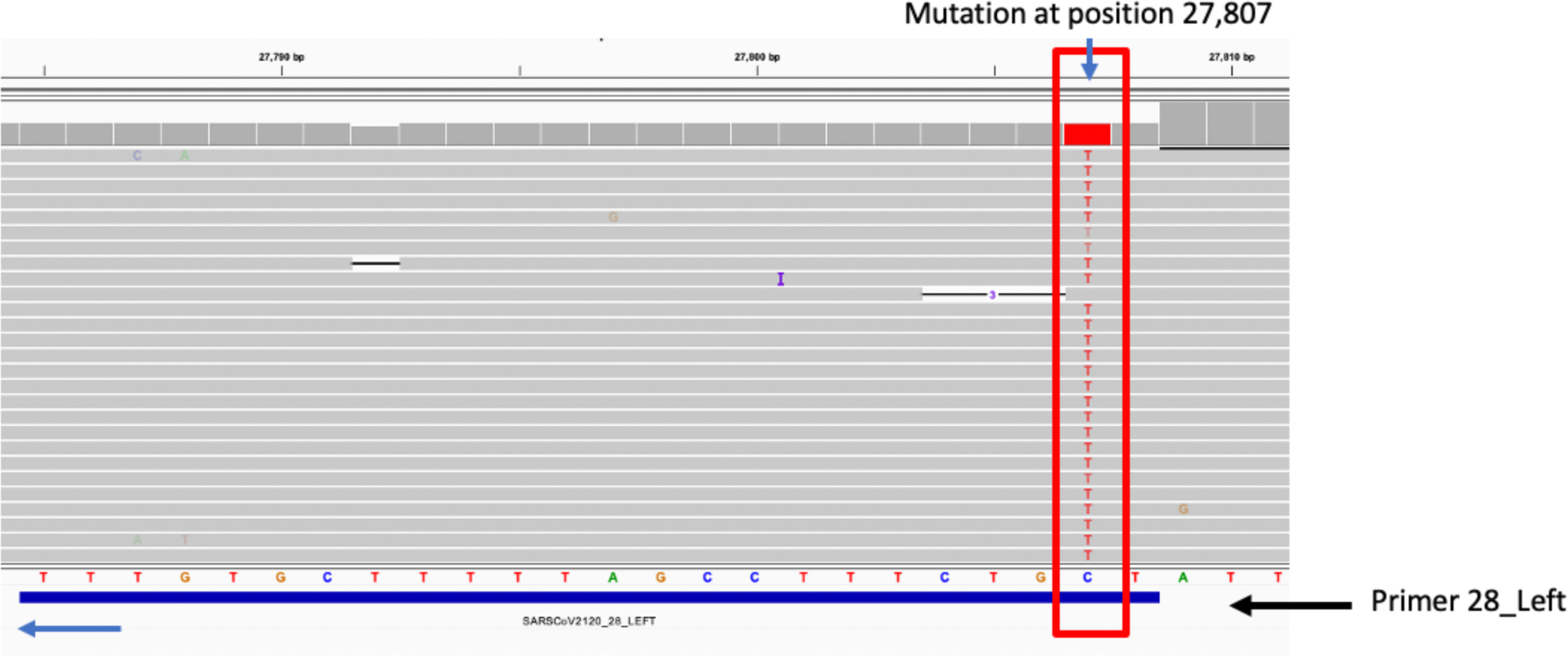
Screenshot of IGV plot showing mutations at position 27,897 of a Delta variant sample sequenced in Nanopore using Midnight primers. This mutation is within the primer binding region for the amplicon 28 (28_LEFT). This is one of the early dropouts observed in most genome sequences generated using Midnight primers.

**Supplementary Figure 2:**
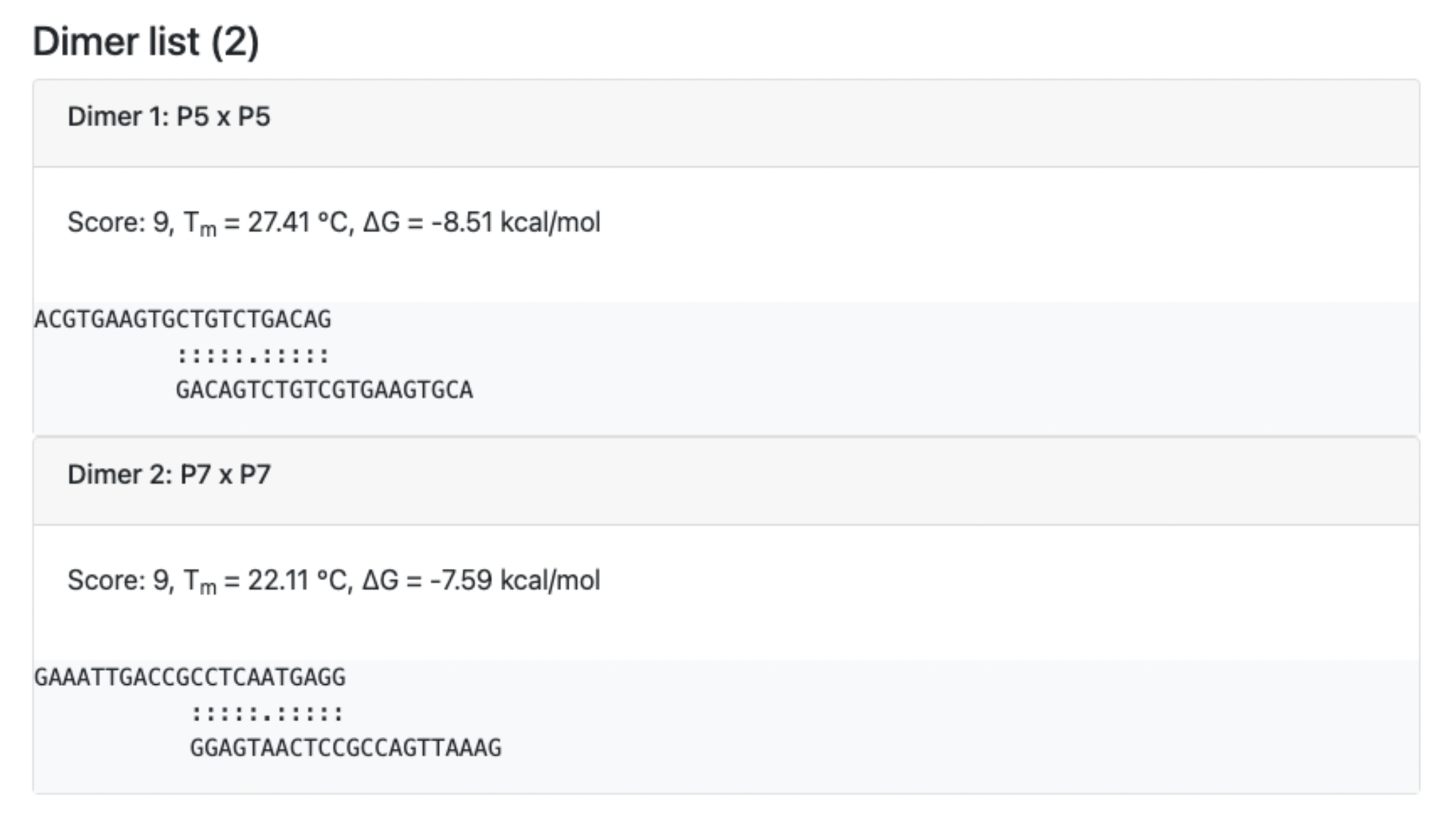
Primer self-interaction for 5_LEFT and 7_LEFT as predicted by MFEprimer.

**Supplementary Figure 3:**
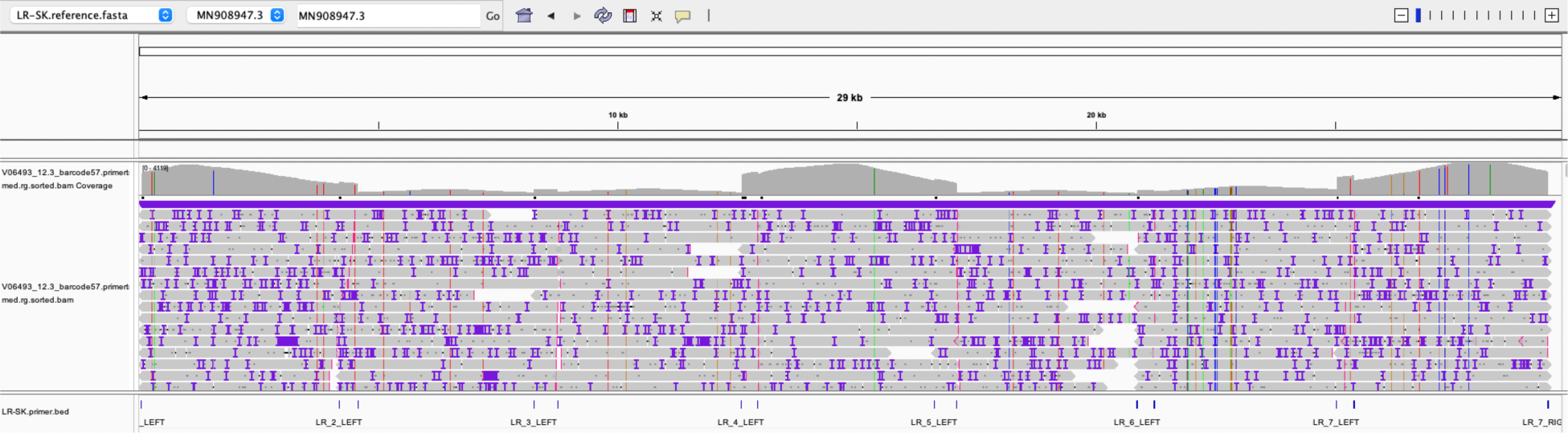
IGV plot showing coverage at different amplicon regions for the sample sequenced using long-range primers. Primers for amplicon regions 2, 3, 5, and 6 were redesigned to increase the coverage at these regions, using reference genomes from GenBank that were collected from December 2022 to March 2023.

**Supplementary Figure 4:**
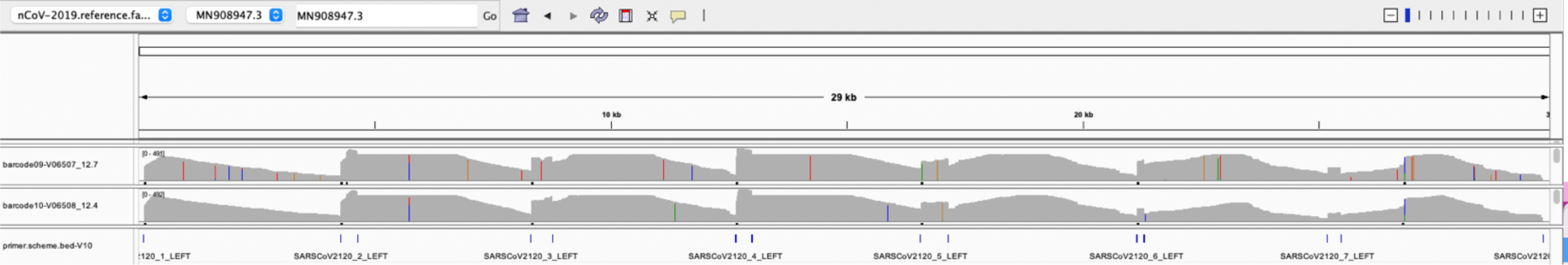
IGV plot showing coverage for three samples with CT values sequenced using updated primer schemes. The samples were accurately assigned a lineage and passed quality.

**Supplementary Table 1:**
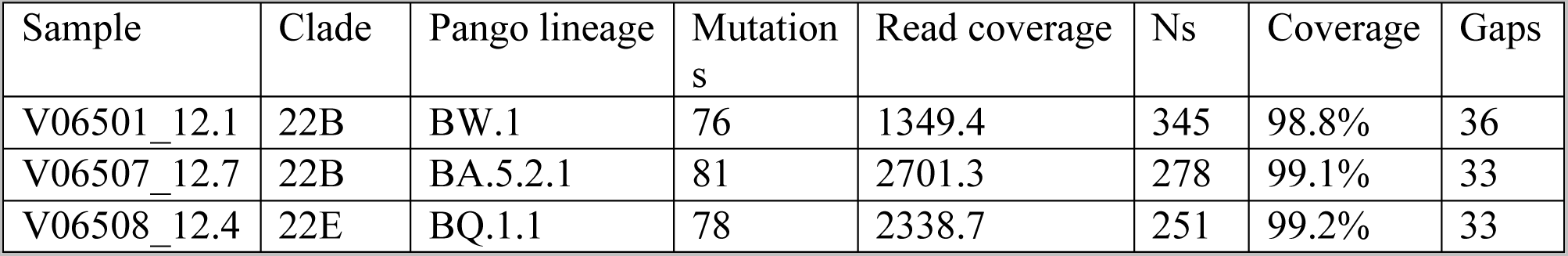
Nextclade results for three samples sequencing using updated long-range primers. The samples had a CT value of 12.

## References

1. Aksamentov, Ivan, Cornelius Roemer, Emma B. Hodcroft, and Richard A. Neher. “Nextclade: Clade Assignment, Mutation Calling and Quality Control for Viral Genomes.” Journal of Open Source Software 6, no. 67 (November 30, 2021): 3773. https://doi.org/10.21105/joss.03773.

2. Alkam, Duah, Piroon Jenjaroenpun, Thidathip Wongsurawat, Zulema Udaondo, Preecha Patumcharoenpol, Michael Robeson, Dirk Haselow, et al. “Genomic Characterization of Mumps Viruses from a Large-Scale Mumps Outbreak in Arkansas, 2016.” *Infection*, Genetics, and Evolution 75 (November 1, 2019): 103965. https://doi.org/10.1016/j.meegid.2019.103965.

3. Arana, Carlos, Chaoying Liang, Matthew Brock, Bo Zhang, Jinchun Zhou, Li Chen, Brandi Cantarel, Jeffrey SoRelle, Lora V. Hooper, and Prithvi Raj. “A Short plus Long-Amplicon Based Sequencing Approach Improves Genomic Coverage and Variant Detection in the SARS-CoV-2 Genome.” PLOS ONE 17, no. 1 (January 13, 2022): e0261014. https://doi.org/10.1371/journal.pone.0261014.

4. Bakhshandeh, Behnaz, Zohreh Jahanafrooz, Ardeshir Abbasi, Matin Babaee Goli, Mahya Sadeghi, Mohammad Sadeq Mottaqi, and Maryam Zamani. “Mutations in SARS-CoV-2; Consequences in Structure, Function, and Pathogenicity of the Virus.” Microbial Pathogenesis 154 (May 1, 2021): 104831. https://doi.org/10.1016/j.micpath.2021.104831.

5. Bei, Yanxia, Kaylinnette Pinet, Kyle B. Vrtis, Janine G. Borgaro, Luo Sun, Matthew Campbell, Lynne Apone, Bradley W. Langhorst, and Nicole M. Nichols. “Overcoming Variant Mutation-Related Impacts on Viral Sequencing and Detection Methodologies.” Frontiers in Medicine 9 (2022). https://www.frontiersin.org/articles/10.3389/fmed.2022.989913.

6. Brito, Anderson F., Elizaveta Semenova, Gytis Dudas, Gabriel W. Hassler, Chaney C. Kalinich, Moritz U. G. Kraemer, Joses Ho, et al. “Global Disparities in SARS-CoV-2 Genomic Surveillance.” Nature Communications 13, no. 1 (November 16, 2022): 7003. https://doi.org/10.1038/s41467-022-33713-y.

7. Butler, Daniel, Christopher Mozsary, Cem Meydan, Jonathan Foox, Joel Rosiene, Alon Shaiber, David Danko, et al. “Shotgun Transcriptome, Spatial Omics, and Isothermal Profiling of SARS-CoV-2 Infection Reveals Unique Host Responses, Viral Diversification, and Drug Interactions.” Nature Communications 12, no. 1 (March 12, 2021): 1660. https://doi.org/10.1038/s41467-021-21361-7.

8. Carabelli, Alessandro M., Thomas P. Peacock, Lucy G. Thorne, William T. Harvey, Joseph Hughes, Thushan I. de Silva, Sharon J. Peacock, et al. “SARS-CoV-2 Variant Biology: Immune Escape, Transmission and Fitness.” *Nature Reviews Microbiology*, January 18, 2023, 1–16. https://doi.org/10.1038/s41579-022-00841-7.

9. Carbo, Ellen C., Igor A. Sidorov, Jessika C. Zevenhoven-Dobbe, Eric J. Snijder, Eric C. Claas, Jeroen F. J. Laros, Aloys C. M. Kroes, and Jutte J. C. de Vries. “Coronavirus Discovery by Metagenomic Sequencing: A Tool for Pandemic Preparedness.” Journal of Clinical Virology 131 (October 1, 2020): 104594. https://doi.org/10.1016/j.jcv.2020.104594.

10. Charre, Caroline, Christophe Ginevra, Marina Sabatier, Hadrien Regue, Grégory Destras, Solenne Brun, Gwendolyne Burfin, et al. “Evaluation of NGS-Based Approaches for SARS-CoV-2 Whole Genome Characterisation.” Virus Evolution 6, no. 2 (July 1, 2020): veaa075. https://doi.org/10.1093/ve/veaa075.

11. Chatterjee, Srijan, Manojit Bhattacharya, Sagnik Nag, Kuldeep Dhama, and Chiranjib Chakraborty. “A Detailed Overview of SARS-CoV-2 Omicron: Its Sub-Variants, Mutations and Pathophysiology, Clinical Characteristics, Immunological Landscape, Immune Escape, and Therapies.” Viruses 15, no. 1 (January 2023): 167. https://doi.org/10.3390/v15010167.

12. Cherif, Emira, Fatou Seck Thiam, Mohammad Salma, Georgina Rivera-Ingraham, Fabienne Justy, Theo Deremarque, Damien Breugnot, Jean-Claude Doudou, Rodolphe Elie Gozlan, and Marine Combe. “ONTdeCIPHER: An Amplicon-Based Nanopore Sequencing Pipeline for Tracking Pathogen Variants.” Bioinformatics 38, no. 7 (March 28, 2022): 2033–35. https://doi.org/10.1093/bioinformatics/btac043.

13. Clark, Cyndi, Joshua Schrecker, Matthew Hardison, and Michael S. Taitel. “Validation of Reduced S-Gene Target Performance and Failure for Rapid Surveillance of SARS-CoV-2 Variants.” PLOS ONE 17, no. 10 (October 3, 2022): e0275150. https://doi.org/10.1371/journal.pone.0275150.

14. Constantinides, Bede, Hermione Webster, Jessica Gentry, Jasmine Bastable, Laura Dunn, Sarah Oakley, Jeremy Swann, et al. “Rapid Turnaround Multiplex Sequencing of SARS-CoV-2: Comparing Tiling Amplicon Protocol Performance.” medRxiv, January 1, 2022. https://doi.org/10.1101/2021.12.28.21268461.

15. Coronaviridae Study Group of the International Committee on Taxonomy of Viruses. “The Species Severe Acute Respiratory Syndrome-Related Coronavirus: Classifying 2019-NCoV and Naming It SARS-CoV-2.” Nature Microbiology 5, no. 4 (April 2020): 536–44. https://doi.org/10.1038/s41564-020-0695-z.

16. Cotten, Matthew, Dan Lule Bugembe, Pontiano Kaleebu, and My V.T. Phan. “Alternate Primers for Whole-Genome SARS-CoV-2 Sequencing.” Virus Evolution 7, no. 1 (January 20, 2021): veab006. https://doi.org/10.1093/ve/veab006.

17. Davies, Nicholas G., Christopher I. Jarvis, W. John Edmunds, Nicholas P. Jewell, Karla Diaz-Ordaz, and Ruth H. Keogh. “Increased Mortality in Community-Tested Cases of SARS-CoV-2 Lineage B.1.1.7.” Nature 593, no. 7858 (May 2021): 270–74. https://doi.org/10.1038/s41586-021-03426-1.

18. Davis, James J., S. Wesley Long, Paul A. Christensen, Randall J. Olsen, Robert Olson, Maulik Shukla, Sishir Subedi, Rick Stevens, and James M. Musser. “Analysis of the ARTIC Version 3 and Version 4 SARS-CoV-2 Primers and Their Impact on the Detection of the G142D Amino Acid Substitution in the Spike Protein.” Microbiology Spectrum 9, no. 3 (December 8, 2021): e01803–21. https://doi.org/10.1128/Spectrum.01803-21.

19. Deng, Xianding, Asmeeta Achari, Scot Federman, Guixia Yu, Sneha Somasekar, Inês Bártolo, Shigeo Yagi, et al. “Metagenomic Sequencing with Spiked Primer Enrichment for Viral Diagnostics and Genomic Surveillance.” Nature Microbiology 5, no. 3 (2020): 443–54. https://doi.org/10.1038/s41564-019-0637-9.

20. Escalera, Alba, Ana S. Gonzalez-Reiche, Sadaf Aslam, Ignacio Mena, Manon Laporte, Rebecca L. Pearl, Andrea Fossati, et al. “Mutations in SARS-CoV-2 Variants of Concern Link to Increased Spike Cleavage and Virus Transmission.” Cell Host & Microbe 30, no. 3 (March 9, 2022): 373–387.e7. https://doi.org/10.1016/j.chom.2022.01.006.

21. Freed, Nikki E, Markéta Vlková, Muhammad B Faisal, and Olin K Silander. “Rapid and Inexpensive Whole-Genome Sequencing of SARS-CoV-2 Using 1200 Bp Tiled Amplicons and Oxford Nanopore Rapid Barcoding.” Biology Methods and Protocols 5, no. 1 (January 1, 2020): bpaa014. https://doi.org/10.1093/biomethods/bpaa014.

22. Galloway, Summer E., Prabasaj Paul, Duncan R. MacCannell, Michael A. Johansson, John T. Brooks, Adam MacNeil, Rachel B. Slayton, et al. “Emergence of SARS-CoV-2 B.1.1.7 Lineage — United States, December 29, 2020–January 12, 2021.” Morbidity and Mortality Weekly Report 70, no. 3 (January 22, 2021): 95–99. https://doi.org/10.15585/mmwr.mm7003e2.

23. Gerber, Zuzana, Christian Daviaud, Damien Delafoy, Florian Sandron, Enagnon Kazali Alidjinou, Jonathan Mercier, Sylvain Gerber, et al. “A Comparison of High-Throughput SARS-CoV-2 Sequencing Methods from Nasopharyngeal Samples.” Scientific Reports 12, no. 1 (July 22, 2022): 12561. https://doi.org/10.1038/s41598-022-16549-w.

24. Grubaugh, Nathan D., Karthik Gangavarapu, Joshua Quick, Nathaniel L. Matteson, Jaqueline Goes De Jesus, Bradley J. Main, Amanda L. Tan, et al. “An Amplicon-Based Sequencing Framework for Accurately Measuring Intrahost Virus Diversity Using PrimalSeq and IVar.” Genome Biology 20, no. 1 (January 8, 2019): 8. https://doi.org/10.1186/s13059-018-1618-7

25. He, Xuemei, Weiqi Hong, Xiangyu Pan, Guangwen Lu, and Xiawei Wei. “SARS-CoV-2 Omicron Variant: Characteristics and Prevention.” MedComm 2, no. 4 (2021): 838–45. https://doi.org/10.1002/mco2.110.

26. Hoffmann, Markus, Prerna Arora, Rüdiger Groß, Alina Seidel, Bojan F. Hörnich, Alexander S. Hahn, Nadine Krüger, et al. “SARS-CoV-2 Variants B.1.351 and P.1 Escape from Neutralizing Antibodies.” Cell 184, no. 9 (April 29, 2021): 2384–2393.e12. https://doi.org/10.1016/j.cell.2021.03.036.

27. Huang, Chaolin, Yeming Wang, Xingwang Li, Lili Ren, Jianping Zhao, Yi Hu, Li Zhang, et al. “Clinical Features of Patients Infected with 2019 Novel Coronavirus in Wuhan, China.” The Lancet 395, no. 10223 (February 15, 2020): 497–506. https://doi.org/10.1016/S0140-6736(20)30183-5.

28. Issa, Elio, Georgi Merhi, Balig Panossian, Tamara Salloum, and Sima Tokajian. “SARS-CoV-2 and ORF3a: Nonsynonymous Mutations, Functional Domains, and Viral Pathogenesis.” MSystems 5, no. 3 (May 5, 2020): e00266–20. https://doi.org/10.1128/mSystems.00266-20.

29. Itokawa, Kentaro, Tsuyoshi Sekizuka, Masanori Hashino, Rina Tanaka, and Makoto Kuroda. “Disentangling Primer Interactions Improves SARS-CoV-2 Genome Sequencing by Multiplex Tiling PCR.” Edited by Ruslan Kalendar. PLOS ONE 15, no. 9 (September 18, 2020): e0239403. https://doi.org/10.1371/journal.pone.0239403.

30. Katoh, Kazutaka, John Rozewicki, and Kazunori D Yamada. “MAFFT Online Service: Multiple Sequence Alignment, Interactive Sequence Choice and Visualization.” Briefings in Bioinformatics 20, no. 4 (July 19, 2019): 1160–66. https://doi.org/10.1093/bib/bbx108.

31. Kuchinski, Kevin S., Jason Nguyen, Tracy D. Lee, Rebecca Hickman, Agatha N. Jassem, Linda M. N. Hoang, Natalie A. Prystajecky, and John R. Tyson. “Mutations in Emerging Variant of Concern Lineages Disrupt Genomic Sequencing of SARS-CoV-2 Clinical Specimens.” International Journal of Infectious Diseases 114 (January 1, 2022): 51–54. https://doi.org/10.1016/j.ijid.2021.10.050.

32. Kupferschmidt, Kai, and Meredith Wadman. “Delta Variant Triggers New Phase in the Pandemic.” Science 372, no. 6549 (June 25, 2021): 1375–76. https://doi.org/10.1126/science.372.6549.1375.

33. Julenius, Karin, and Anders Gorm Pedersen. “Protein Evolution Is Faster Outside the Cell.” Molecular Biology and Evolution 23, no. 11 (November 2006): 2039–48. https://doi.org/10.1093/molbev/msl081.

34. Lambisia, Arnold W., Khadija S. Mohammed, Timothy O. Makori, Leonard Ndwiga, Maureen W. Mburu, John M. Morobe, Edidah O. Moraa, et al. “Optimization of the SARS-CoV-2 ARTIC Network V4 Primers and Whole Genome Sequencing Protocol.” Frontiers in Medicine 9 (February 17, 2022): 836728. https://doi.org/10.3389/fmed.2022.836728.

35. Li, Heng, Bob Handsaker, Alec Wysoker, Tim Fennell, Jue Ruan, Nils Homer, Gabor Marth, Goncalo Abecasis, Richard Durbin, and 1000 Genome Project Data Processing Subgroup. “The Sequence Alignment/Map Format and SAMtools.” Bioinformatics 25, no. 16 (August 15, 2009): 2078–79. https://doi.org/10.1093/bioinformatics/btp352.

36. Liu, Tiantian, Zhong Chen, Wanqiu Chen, Xin Chen, Maryam Hosseini, Zhaowei Yang, Jing Li, et al. “A Benchmarking Study of SARS-CoV-2 Whole-Genome Sequencing Protocols Using COVID-19 Patient Samples.” IScience 24, no. 8 (August 20, 2021): 102892. https://doi.org/10.1016/j.isci.2021.102892.

37. Madhi, Shabir A., Vicky Baillie, Clare L. Cutland, Merryn Voysey, Anthonet L. Koen, Lee Fairlie, Sherman D. Padayachee, et al. ”Efficacy of the ChAdOx1 NCoV-19 Covid-19 Vaccine against the B.1.351 Variant.” *The New England Journal of Medicine*, March 16, 2021, NEJMoa2102214. https://doi.org/10.1056/NEJMoa2102214.

38. Meng, Bo, Steven A. Kemp, Guido Papa, Rawlings Datir, Isabella A. T. M. Ferreira, Sara Marelli, William T. Harvey, et al. “Recurrent Emergence of SARS-CoV-2 Spike Deletion H69/V70 and Its Role in the Alpha Variant B.1.1.7.” Cell Reports 35, no. 13 (June 29, 2021). https://doi.org/10.1016/j.celrep.2021.109292.

39. O’Toole, Áine, Emily Scher, Anthony Underwood, Ben Jackson, Verity Hill, John T McCrone, Rachel Colquhoun, et al. “Assignment of Epidemiological Lineages in an Emerging Pandemic Using the Pangolin Tool.” Virus Evolution 7, no. 2 (December 1, 2021): veab064. https://doi.org/10.1093/ve/veab064.

40. Quick, Josh. ”NCoV-2019 Sequencing Protocol,” January 22, 2020. https://www.protocols.io/view/ncov-2019-sequencing-protocol-bbmuik6w.

41. Quick, Joshua, Nathan D. Grubaugh, Steven T. Pullan, Ingra M. Claro, Andrew D. Smith, Karthik Gangavarapu, Glenn Oliveira, et al. “Multiplex PCR Method for MinION and Illumina Sequencing of Zika and Other Virus Genomes Directly from Clinical Samples.” Nature Protocols 12, no. 6 (June 2017): 1261–76. https://doi.org/10.1038/nprot.2017.066.

42. Rehn, Alexandra, Peter Braun, Mandy Knüpfer, Roman Wölfel, Markus H. Antwerpen, and Mathias C. Walter. “Catching SARS-CoV-2 by Sequence Hybridization: A Comparative Analysis.” MSystems 6, no. 4 (August 3, 2021): e00392–21. https://doi.org/10.1128/mSystems.00392-21.

43. Sanderson, Theo, and Jeffrey C. Barrett. “Variation at Spike Position 142 in SARS-CoV-2 Delta Genomes Is a Technical Artifact Caused by Dropout of a Sequencing Amplicon.” Wellcome Open Research 6 (November 10, 2021): 305. https://doi.org/10.12688/wellcomeopenres.17295.1.

44. Thaweethai, Tanayott, Sarah E. Jolley, Elizabeth W. Karlson, Emily B. Levitan, Bruce Levy, Grace A. McComsey, Lisa McCorkell, et al. ”Development of a Definition of Postacute Sequelae of SARS-CoV-2 Infection.” *JAMA*, May 25, 2023. https://doi.org/10.1001/jama.2023.8823.

45. Tyson, John R, Phillip James, David Stoddart, Natalie Sparks, Arthur Wickenhagen, Grant Hall, Ji Hyun Choi, et al. ”Improvements to the ARTIC Multiplex PCR Method for SARS-CoV-2 Genome Sequencing Using Nanopore.” *BioRxiv*, September 4, 2020, 2020.09.04.283077. https://doi.org/10.1101/2020.09.04.283077.

46. Vacca, Davide, Antonino Fiannaca, Fabio Tramuto, Valeria Cancila, Laura La Paglia, Walter Mazzucco, Alessandro Gulino, et al. “Direct RNA Nanopore Sequencing of SARS-CoV-2 Extracted from Critical Material from Swabs.” *Life (Basel*, Switzerland) 12, no. 1 (January 4, 2022): 69. https://doi.org/10.3390/life12010069.

47. Wang, Kun, Haiwei Li, Yue Xu, Qianzhi Shao, Jianming Yi, Ruichao Wang, Wanshi Cai, et al. “MFEprimer-3.0: Quality Control for PCR Primers.” Nucleic Acids Research 47, no. W1 (July 2, 2019): W610–13. https://doi.org/10.1093/nar/gkz351.

48. Wassenaar, Trudy M, Visanu Wanchai, Gregory Buzard, and David W Ussery. “The First Three Waves of the Covid-19 Pandemic Hint at a Limited Genetic Repertoire for SARS-CoV-2.” FEMS Microbiology Reviews 46, no. 3 (May 1, 2022): fuac003. https://doi.org/10.1093/femsre/fuac003.

49. Wick, Ryan R., Louise M. Judd, and Kathryn E. Holt. “Performance of Neural Network Basecalling Tools for Oxford Nanopore Sequencing.” Genome Biology 20, no. 1 (June 24, 2019): 129. https://doi.org/10.1186/s13059-019-1727-y.

50. Wu, Fan, Su Zhao, Bin Yu, Yan-Mei Chen, Wen Wang, Zhi-Gang Song, Yi Hu, et al. “A New Coronavirus Associated with Human Respiratory Disease in China.” Nature 579, no. 7798 (March 2020): 265–69. https://doi.org/10.1038/s41586-020-2008-3.

51. Xiao, Minfeng, Xiaoqing Liu, Jingkai Ji, Min Li, Jiandong Li, Lin Yang, Wanying Sun, et al. “Multiple Approaches for Massively Parallel Sequencing of SARS-CoV-2 Genomes Directly from Clinical Samples.” Genome Medicine 12, no. 1 (June 30, 2020): 57. https://doi.org/10.1186/s13073-020-00751-4.

52. Zhou, Peng, Xing-Lou Yang, Xian-Guang Wang, Ben Hu, Lei Zhang, Wei Zhang, Hao-Rui Si, et al. “A Pneumonia Outbreak Associated with a New Coronavirus of Probable Bat Origin.” Nature 579, no. 7798 (March 2020): 270–73. https://doi.org/10.1038/s41586-020-2012-7.

